# High-Resolution EEG Source Reconstruction from PCA-Corrected BEM-FMM Reciprocal Basis Funcions: A Study with Visual Evoked Potentials from Intermittent Photic Stimulation

**DOI:** 10.1101/2025.07.11.664246

**Authors:** Guillermo Nuñez Ponasso, Derek A. Drumm, Hannes Oppermann, Abbie Wang, Gregory M. Noetscher, Burkhard Maess, Thomas R. Knösche, Sergey N. Makaroff, Jens Haueisen

## Abstract

Modern automated human head segmentations can generate high-resolution computational meshes involving many non-nested tissues. However, most source reconstruction software is limited to 3 –4 nested layers of low resolution and a small number of dipolar sources∼ 10, 000.

Recently, we introduced modeling techniques for source reconstruction of magnetoencephalographic (MEG) signals using the reciprocal approach and the boundary element fast multipole method (BEM-FMM). The technique of BEM-FMM can process both nested and non-nested models with as many as 4 million surface elements.

In this paper, we present an analogue technique for source reconstruction of electroencephalographic (EEG) signals based on cortical global basis functions. The present work uses Helmholtz reciprocity to relate the reciprocally-generated lead-field matrices to their direct counterpart, while resolving the issue of possible biases toward the reference electrode.

Our methodology is tested with experimental EEG data collected from a cohort of 12, young and healthy, volunteers subjected to intermittent photic stimulation (IPS). Our novel high-resolution source reconstruction models can have impact on mental health screening as well as brain-computer interfaces.

## 1 Introduction

Source localization, also known as source estimation or source reconstruction, (Knösche and Haueisen 2022) consists of estimating the distribution of neural activity on the cortex that best explains a given set of neurophysiological recordings, either from electroencephalography (EEG), magnetoencephalography (MEG), or other modalities. EEG constitutes the most affordable and widely-available modality of recordings; as such, source localization based on EEG is of great importance for both clinical and research-focused applications of neuroscience, such as mental health screening among many others.

Modern automated segmentations from magnetic resonance imaging (MRI) can generate high resolution triangulated mesh models distinguishing up to 40 human *non-nested* head tissues (IT’IS Foundation 2024); as well as high-resolution *nested* and watertight meshes modeling 5 major human head volumes: skin, skull, cerebrospinal fluid (CSF), gray matter (GM), and white matter (WM), (Fischl et al. 2004; Nielsen et al. 2018). Major source localization packages like the MNE Software (Gramfort et al. 2013; Gramfort et al. 2014), FieldTrip (Oostenveld et al. 2011), or Brainstorm (Tadel et al. 2011) make use of either the boundary element method (BEM) or the finite element method (FEM). In both cases, the solvers used are direct: this means that the EEG/MEG sensor outputs are simulated from specified dipole locations and moments inside the head volume. These solvers are based on constructing and then inverting the so-called system matrix, which depends only on the anatomical models extracted from the MRI segmentation and on a choice of conductivity values for each tissue.

FEM solvers like DUNEuro (Schrader et al. 2021), have the ability to model EEG and MEG with the 5-layer nested models mentioned before. This is because the FEM system matrix is sparse, and solving a sparse system —using, for example, the conjugate-gradient method (Hestenes and Stiefel 1952)— can be achieved in 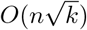 operations, where *n* is the number of volumetric elements and *k* is the matrix’s condition number. On the other hand, BEM solvers (Kybic et al. 2005; Gramfort et al. 2010) are incapable of processing the high-resolution 5-layer meshes because the associated system matrices are dense, and solving linear systems with them involves *O*(*n*^3^) operations, where *n* is the number of surface elements. The main disadvantage of FEM over BEM is that FEM requires volumetric elements; it is therefore harder to model complicated geometries and thin tissue interfaces with FEM, which makes the number of elements needed is much higher. Additionally, the solutions of FEM can only be sampled at the elements of the volumetric meshes, whereas the solutions of BEM can be sampled anywhere in space.

The boundary-element fast multipole method (BEM-FMM) (Makarov et al. 2018; Makarov et al. 2020; Makarov et al. 2021a; Makarov et al. 2021b) is an iterative method that serves as a powerful alternative to FEM and classical BEM. While classical BEM approaches to EEG/MEG modeling use the surface potential as the fundamental unknown (Geselowitz 1967; Geselowitz 1970), the BEM-FMM uses the charge-based formulation (Gelernter and Swihart 1964; Barnard et al. 1967; Makarov et al. 2018; Nuñez Ponasso 2024) coupled with the fast multipole method (FMM) (Greengard and Rokhlin 1987). As an iterative method, the BEM-FMM does not create a system matrix, so it is able to compute fast solutions using highly complex and realistic meshes such as the 40-tissue head models or the highresolution 5-layer head models. In fact, it was found that the BEM-FMM provided the highest accuracy for transcranial magnetic stimulation (TMS) modeling (within computational limitations) among a wide range of methods (Gomez et al. 2020). However, directly computing the EEG/MEG outputs for each possible cortical dipole location is prohibitive for BEM-FMM since the entire computation must be repeated for each dipole.

A solution to this limitation of BEM-FMM is to use the reciprocal approach to the EEG/MEG forward problem. In Nuñez Ponasso et al. 2025 and Drumm et al. 2025, we introduced and tested reciprocal BEM-FMM source reconstruction methods for MEG using high-resolution meshes and significantly improved the accuracy and focality of localizations over previous methodologies. The reciprocal approach is based on the computation of *cortical global basis functions*: for MEG these basis functions consist of the electric field on the cortical surface induced by TMS where a fictitious current is injected through each of the MEG pick-up coils. The reciprocity theorem identifies the cortical global basis functions as the rows of the lead-field matrix (LFM), also known as gain matrix. The reciprocal approach can be then understood as filling-in the LFM by rows, whereas the direct approach fills-in the LFM by columns. Since the LFM has size *M*× *N* (or *M*× 3*N* for freely oriented dipoles (Lin et al. 2006)), where *M* is the number of sensors and *N* is the number of sources in the model, BEM-FMM can easily compute the LFM reciprocally using as many sources as there are elements on the chosen cortical surface (WM, GM, or any surface in between WM and GM). For example, the high-resolution 5-layer nested models of the *headreco* SimNIBS pipeline (Nielsen et al. 2018) have approximately 250,000 triangle elements in the WM mesh; in contrast, the largest number of sources MNE-Python (Gramfort et al. 2013) can process is ∼ 20,000.

Similar to the reciprocal BEM-FMM source reconstruction methods for MEG, we introduce here an analogous methodology for reconstructing EEG sources. There are two major challenges in the reciprocal calculation of EEG source estimates (not present in the MEG equivalent) that we fully address here:

i. While MEG measurements are absolute, all EEG measurements are given relative to a reference electrode. Using the same reference electrode for the computation of all reciprocal basis functions will steer the ensuing localizations towards the reference —from the Bayesian perspective (Knösche and Haueisen 2022) this is equivalent to a poorly-chosen prior distribution. Previous attempts at reciprocal EEG tried to address this issue by optimizing the selection of electrode pairs (Wartman et al. 2022). This selection procedure can be interpreted as a regularization technique for the LFM; however, an explicit linear relationship between the direct LFM and the LFM generated in Wartman et al. 2022 was not established. As such, the applicability of classical source reconstruction methods, like minimum norm estimation (MNE) (Hämäläinen and Ilmoniemi 1994) or dynamic statistical parametric mappings (dSPM) (Dale et al. 2000) was limited. We solve this issue in full by using a single reference electrode, which establishes a one-to-one correspondence between basis functions and EEG channels. By Helmholtz reciprocity, we relate the reciprocal LFM to the direct LFM. Applying a principal component analysis (PCA) correction, we remove potential biases near the reference electrode. After all these procedures, our new reciprocal method can be directly coupled with all classical source reconstruction techniques for EEG.
ii. Reciprocal MEG entails the calculation of forward TMS solutions (with a fictitious current applied to the MEG sensors), whereas reciprocal EEG requires the computation of transcranial electric stimulation (TES) solutions (with a fictitious current injected into the voltage electrodes). The TMS computation involves coils that are well-separated from the head tissues, so the solution of the forward problem is numerically-stable (Weise et al. 2022); on the other hand, the impressed currents of the TES computation lead to singularities directly above the skin, and the method of adaptive mesh refinement (AMR) (Wartman et al. 2024) is needed in order to obtain accurate numerical results. We used the BEM-FMM solver with the AMR of (Wartman et al. 2024) to generate physically-accurate TES solutions.

To test our methodology, we performed source reconstruction of steady-state visual evoked potentials (ssVEP). These ssVEPs describe electrophysiological responses to repetitive visual stimulations at certain frequencies. The characteristic response of neural oscillations to this kind of stimulus was first shown by Adrian and Matthes in the 1930s (Compston 2009). Since then, there has been considerable interest in the study and analysis of ssVEPs in healthy and pathological brains. Cognitive and clinical research is performed with ssVEP experiments, for example, to examine vision (Norcia et al. 2015), attention and working memory (Chota et al. 2024), or diseases like schizophrenia (Schielke and Krekelberg 2022), among others. In the contemporary era, ssVEP-based brain-computer interfaces (BCI) represent a highly prominent domain of research. The most well-known of these BCIs are speller systems. They enable communication between restricted patients (e.g., locked-in patients) and their environment (Li et al. 2021). Systems are also used for the control of external devices, such as a robotic arm (Ai et al. 2023; Zhu et al. 2020), or a wheelchair (Li et al. 2013). In addition to most ssVEP-based BCIs, which perform a pure sensor space analysis, there are studies that already show the added value of source reconstruction (Zarei and Mohammadzadeh Asl 2022; Oikonomou and Kompatsiaris 2020). This can increase the accuracy of the classification, making the systems even more robust against noise and errors and thus providing a direct benefit for the application of the affected persons.

In Section 2, we describe the experimental data collection as well as our methodology for the generation of the basis functions used for source localization. In Section 3 we give an account of source localization results with relevant figures. In Section 4 we give interpret our results highlighting pros and cons, as well as possible implications to BCIs.

## 2 Materials and Methods

### 2.1 Experimental data collection

#### 2.1.1 Study paradigm

We conducted a photic driving experiment using individualized intermittent photic stimulation (IPS) with stimulation frequencies set to half of the individual alpha peak frequency (iAPF) of each volunteer. The stimulation was divided into 20 trials, with 80 flashes per trial (50% duty cycle) and a 5-second break between consecutive trials. Hence, a total of 1600 single stimuli were provided. Twelve healthy volunteers (5 female, 10 right-handed) with a mean age of 29±7 years participated in the study. The Ethics Committee of the Medical Faculty of Friedrich-Schiller University Jena approved the study, and all volunteers gave their written informed consent to the study and the use of the recorded data.

In a first session, we recorded 5 minutes of resting-state EEG to determine the iAPF. Volunteers were seated in a relaxed position with their eyes closed. A 64-channel EEG cap (Ag/AgCl electrodes, waveguardTM original CA-212.s1, ANT Neuro b.v., Hengelo, Netherlands) and a mobile 64-channel referential DC-EEG amplifier (eegoTM amplifier EE-225, ANT Neuro, b.v., Hengelo, Netherlands) were used for the recordings. A central electrode (5Z) served as an online reference, the electrical ground was placed on the left earlobe, and the sampling rate was set to 2048 Hz.

The second session comprised a combined EEG/MEG experiment with the individualized IPS, whereby we report here only on the EEG recordings. Volunteers were seated in a relaxed position with their eyes closed in a magnetically shielded room. EEG during stimulation was acquired employing a MEG-compatible cap comprising 61 Ag/AgCl electrodes (waveguard™ CA-169, ANT b.v., The Netherlands), placed according to the extended 10-20 system, with an online reference at the position AFz, the electrical ground on the left mastoid, and a sampling rate of 1000 Hz. The EEG amplifier provided by a MEG system (MEGIN, Helsinki, Finland) was used for the recordings. We used a Polhemus 3D digitizer to obtain the anatomical landmarks (nasion and both preauricular points), the individual locations of the EEG electrodes, and randomly distributed points over the head surface.

T1-weighted MRIs from all volunteers were acquired using a 3T Magneto Prisma MRI scanner with a 0.8 mm resolution (Siemens Healthineers AG, Forchheim, Germany).

#### 2.1.2 Signal processing

##### Software

Python (Version 3.11.6), Matlab (The Math-Works, Natick, MA, United States, Version R2023b), the library FMM3D (The Flatiron Institute 2012), and the MNE-Python toolbox (Gramfort et al. 2013) within customized scripts were used for all (pre-)processing steps of the EEG recordings.

##### Individual alpha peak frequency

The resting-state EEG recording was only used for the determination of the iAPF. After bandpass filtering of the data (2-45 Hz, Butterworth, 4th order), we visually identified and removed any bad channels, before re-referencing the data to common average reference (CAR). We reduced the number of channels to eight occipital channels and divided the signal into non-overlapping 30-second epochs. The first and last epochs were then removed. Subsequently, we calculated the power spectral density (PSD), using the Welch method (Welch 1967) for each occipital channel and segment. We applied the Savitzky-Golay smoothing filter (Savitzky and Golay 1964) and determined the iAPF as the main peak in the alpha band (8-13 Hz) from the mean across the eight channels and segments.

##### IPS data

We filtered the EEG recordings with a finite-impulse response (FIR) bandpass filter between 2 and 45 Hz for visual inspection and removal of bad channels and bad trials, i.e. flat lines, high amplitudes, and channels with repeating artifacts. Subsequently, we applied independent component analysis (ICA) to detect and reduce the impact of electrocardiogram (ECG) and eye-movement artifacts. The cleaned EEG data were re-referenced to CAR. For further source localization tasks, the recordings were divided into epochs. First, stimulation epochs were created, containing 250 ms of signal, starting 50 ms prior to the onset of each stimulation. Depending on the number of removed trials, approximately 1500 epochs were averaged and used for the subsequent source reconstruction. Second, baseline (non-stimulation, resting state) epochs were created. Epochs were taken from the breaks between each trial and lasted 3.5 seconds. A maximum of 20 epochs was extracted from the data.

### 2.2 Head modeling

MRI T1-weighed data for all 12 healthy volunteers were automatically segmented using the *headreco* routine (Nielsen et al. 2018) of the SimNIBS pipeline (Saturnino et al. 2019). This segmentation provides 5 main compartments: skin, skull, cerebrospinal fluid (CSF), gray matter (GM), and white matter (WM); with two additional (much smaller and less relevant from a modeling standpoint) tissue meshes for the eyes, and ventricles.

### 2.3 Main Concepts in Source Reconstruction and the Reciprocal Approach

To illustrate the key concepts, consider the following simplified description of the EEG forward problem: Suppose we have mass neural activation at a single cortical location **p**; this type of activation may be modeled using a current dipole (either finite length or point dipole) centered at **p** with a given dipole moment **q** (Nunez 1981; Hämäläinen et al. 1993). At the same time assume that we have a pair of electrodes *e*_1_ and *e*_0_ which measure the difference in skin potential caused by the mass neuronal activation; this is an EEG set-up with one single channel given by *e*_1_ and one reference electrode given by *e*_0_. We denote by **r**_1_ and **r**_0_ be the locations on the skin surface of the electrodes *e*_1_ and *e*_0_ respectively. The task of the forward EEG problem is to determine the difference of potentials

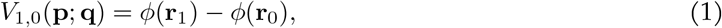

In bioelectromagnetism, the quasi-static approximation to Maxwell’s equations is justified (Plonsey and Heppner 1967). Therefore, by linearity, the more general situation where we have several EEG sensors, and where we may have neural activation distributed over larger cortical regions can be reduced to the computation of the potential differences *V*_*i*,0_(**p**; **q**) for the entire collection of sensors *e*_*i*_ in reference to electrode *e*_0_ and for all possible position/moment pairs (**p, q**).

To perform EEG source localization, one prescribes a finite collection of possible position/moment pairs *S*, known as the *source space*, and computes the so-called *leadfield matrix* (or *gain matrix*) *L* whose rows are indexed by (non-reference) electrode sensors and whose columns are indexed by locations **p**_*j*_, and whose (*i, j*) entry is given by

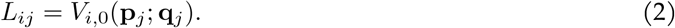

The leadfield matrix is used to construct an *inverse operator* (see §2.7) which provides an estimation of the activation regions that best explain the experimentally measured EEG data. In this manner, the relationship between sources and measurements is established by the leadfield matrix.

In the direct approach, one computes the values *V*_*i*,0_(**p**_*j*_; **q**_*j*_) for each *j* one by one; in other words, the leadfield matrix is “filled in by columns”. This approach is the most common, and in most source-localization software packages the computation is done using the potential-based formulation of the *boundary element method* (BEM) (Geselowitz 1967; Geselowitz 1970). This approach is computationally expensive and, given that the BEM system matrix is dense, the number of tissue layers typically becomes limited to 3-layers (skin, outer skull, and inner skull), their spatial resolution must be reduced, and the number of sources we can employ becomes limited to ∼ 10,000—20,000.

In the reciprocal approach, we use Helmholtz’s reciprocity relation (Helmholtz 1853; Plonsey 1963) to compute the potential differences *V*_*i*,0_(**p**_*j*_, **q**_*j*_) for the entire collection of positions and moments (**p**_*j*_, **q**_*j*_) all at once. In other words, reciprocity allows us to “fill-in the leadfield matrix by rows”. Helmholt’z Reciprocity (see §2.4) establishes that *V*_*i*,0_(**p**_*j*_, **q**_*j*_) can be computed as the electric field at position **p**_*j*_ upon injection of an outflowing current in the reference electrode *e*_0_ and an inflowing current in the electrode *e*_*i*_.

### 2.4 Helmholtz Reciprocity for EEG dipoles and Modeling Aspects

We give the statement of Helmholtz’s reciprocity relation relating EEG dipoles and TES using current electrodes. The reader should note that this does not limit the application of reciprocity to EEG systems based on current electrodes: reciprocity can be thought as establishing a relationship between two modeling problems: on the one hand computing voltage differences between two on-skin locations caused by an EEG dipole and, on the other hand, computing the electric field at the dipole location caused by a pair of current electrodes placed in those same skin locations.

Consider a current point-dipole at position **p** and moment **q** = *q***d**, where *q* = 1 [A] is the magnitude of the dipole current and **d** is the direction component of the moment, measured in [m]. Denote by *ϕ*_1_ the electric potential on the skin caused by the current dipole. Consider two current electrodes *e*_*i*_ and *e*_0_ centered at skin positions **r**_*i*_ and **r**_0_, respectively. Inject a total current of 1 [A] flowing into the skin from electrode *e*_*i*_ and a current of 1 [A] flowing out of the skin from electrode *e*_0_. Denote by **E**_2_(**r**; *e*_*i*_) the electric field elicited by this TES arrangement with electrode pair (*e*_*i*_, *e*_0_) on the cortical location **r**. Then, Helmholtz Reciprocity states (cf. Plonsey 1963):

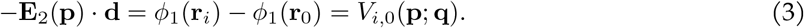

For a derivation of this reciprocity relation based on the quasi-static approximation to Maxwell’s equations see Appendix A. The equation above is a simplification for point electrodes. While the point electrode model is sometimes used in practice (Knösche and Haueisen 2022), we note that a version of this equation for surface electrodes can be derived similarly (although with more technical details). Appendix A also contains an alternative derivation based on circuital reciprocity.

By Equation 2, the computation of the TES solution for the electrode pair (*e*_*i*_, *e*_0_) gives the *i*-th row of the leadfield matrix.

Coupling EEG reciprocity with BEM-FMM has two main advantages: 1) the computation of TES solutions with current electrodes involves only Fredholm integral equations of the second kind (Makarov et al. 2021a; Nuñez Ponasso 2024); and 2) restricting to dipoles oriented normally to the cortical surface (Hämäläinen et al. 1993), the term **E**_2_(**p**) ·**d** is equal to the charge density at **p**, so hence the computation of volume fields is not necessary.

From a modeling perspective, certain approaches like the *finite element method* (FEM), or the potential-based BEM, are much more suited to computing TES solutions for voltage electrodes. This is because the fundamental unknown in these methods is the electric potential. For BEM-FMM, the fundamental unknown is the charge density, therefore the boundary condition for voltage electrodes takes the shape of a Fredholm integral equation of the first kind (which are more challenging to solve). This being said, using a calibration approach, we can model voltage electrodes using current electrodes and vice-versa: for example, in the first case it is sufficient to adjust the impressed current in each electrode until the desired impressed voltage is reached.

### 2.5 Computation of forward solutions for TES with voltage electrodes

To bridge our BEM-FMM reciprocal approach with the FEM and BEM techniques, we will model the TES stimulation using voltage electrodes. We used the solver described in Wartman et al. 2024, which is based on the BEM-FMM solver for voltage electrodes described in Makarov et al. 2021a. This approach incorporates AMR, which is an automated procedure to locally refine the model’s computational meshes wherever the discretization error is highest.

A 61-channel EEG set-up (described in §2.1) was imprinted onto the triangulated skin mesh created from each volunteer, see Figure 1. A total of 60 basis functions were created by simulating the activation of the reference electrode as a fixed cathode (−1 [mA]) and every channel electrode as an anode (+1 [mA]). We refer to each of the TES solutions for electrode pair (*e*_*i*_, *e*_0_) as the *i*-th reciprocal basis function, where *i* = 1, 2, …, 60. By Equations 2 and 3 this corresponds precisely to the *i*-th row of the leadfield matrix. Examples of basis functions can be visualized in Figure 2; observe that there is a strong charge deposition near the reference electrode on all displayed basis functions.

**Figure 1.**
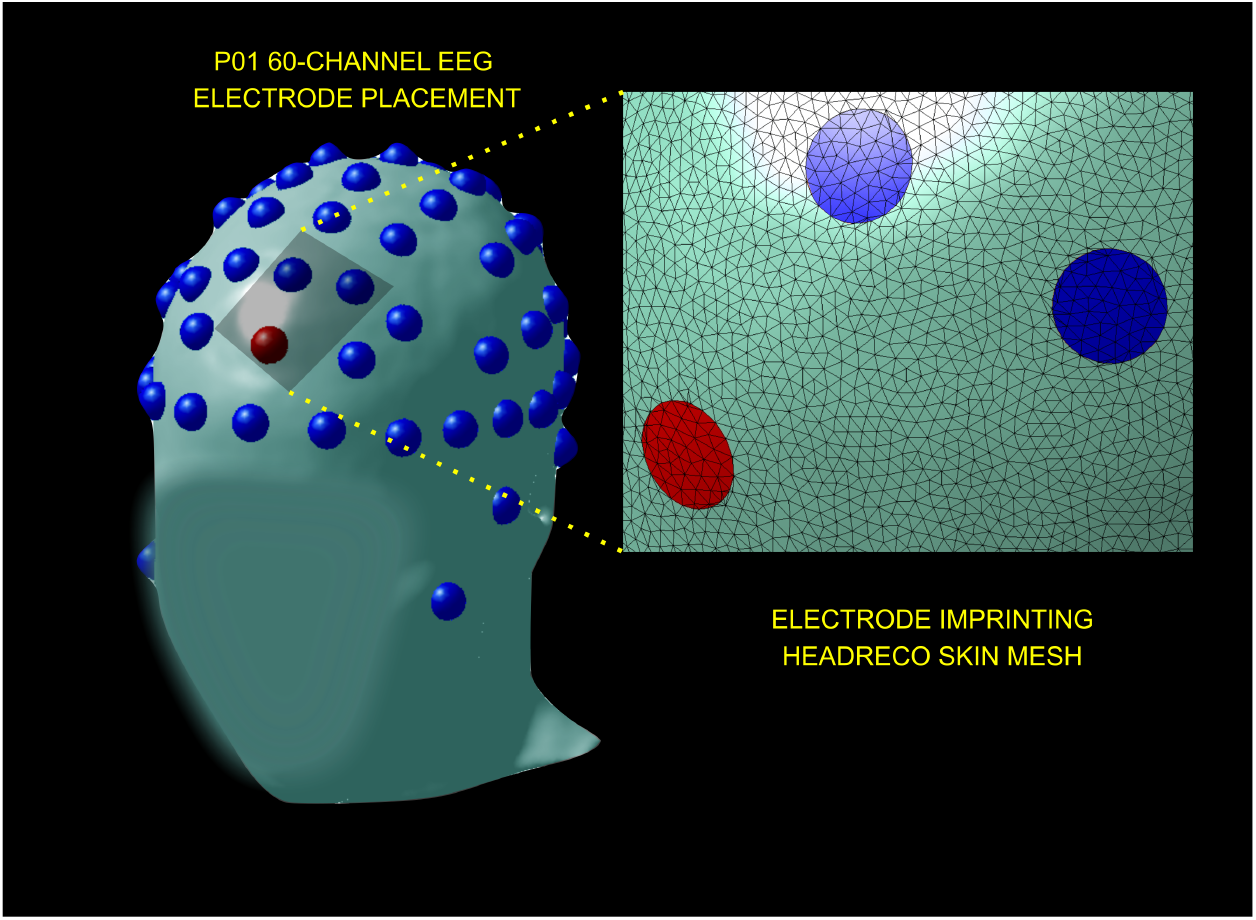
61-channel EEG system placement on volunteer P01 (facial features are blurred for the purpose of privacy). The reference electrode is indicated with a red sphere. After co-registration of electrodes, these are imprinted into the model by refining the skin mesh so as to represent a circular contact surface accurately.

**Figure 2.**
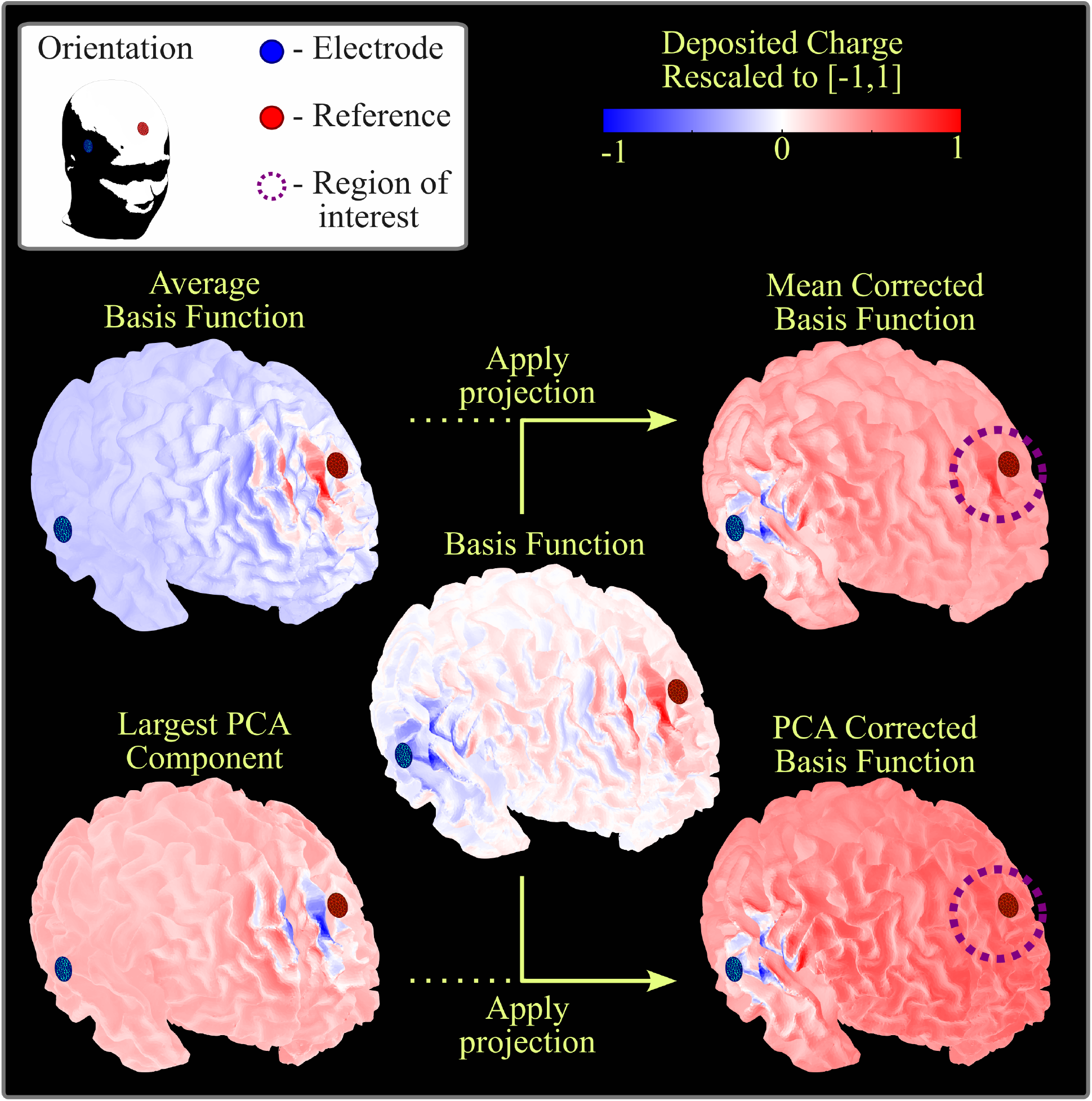
An example of the effect of applying projections to a basis function (central image). Here we show a basis function for volunteer P01, with the EEG electrode in blue and the reference electrode in red. The basis function has clear peaks near the EEG and reference electrodes. Notice how the average basis function (upper-left) —i.e., the average over the rows of the leadfield matrix— has the largest magnitude near the reference electrode. After subtracting this average to obtain the mean corrected basis function (upper-right), there is still a high magnitude near the reference (purple dashed circle). In comparison, when projecting to the space orhtogonal to the largest PCA component of the leadfield matrix (lower-left), the peak in potential near the reference is removed (lower-right).

### 2.6 Processing of basis functions with PCA

To avoid biasing the source reconstructions toward the reference electrode, we remove the component which is present in all of the basis functions. Removing the average is insufficient, as Figure 2 shows. The reason for this is that although there is positive charge near the reference in all basis functions, this charge distribution is not identical in all cases. A better way to determine the common component among all basis functions is to apply a dimensionality reduction via principal component analysis (PCA). Namely, we will project all basis functions onto the space orthogonal to the largest principal component. In Figure 2, we can observe how the charge near the reference electrode has completely disappeared in the PCA-corrected basis functions.

#### Why does projecting onto the space orthogonal to the largest principal component remove the charge distribution near the reference electrode?

From a statistical point of view, the presence of a persistent charge distribution near the reference is reflected as a high correlation between all basis functions. Therefore, the covariance matrix of the basis functions —which is given by the 60×60 matrix *LL*^T^, where *L* is the lead-field matrix— is a matrix with positive off-diagonal entries; i.e. the correlation matrix will have off diagonal entries close to 1. This implies that the eigenvector corresponding to the largest eigenvalue (i.e. the largest principal component) will be close to the all-ones vector. In fact it would be equal to the all-ones vector if and only if removing the average completely removes the bias to the reference electrode. The mathematical justification for these claims is given by Perron-Frobenius theory (Perron 1907; Frobenius 1912), but a formal discussion of this is out of the scope of this paper.

### 2.7 Source Reconstruction

We manually identified the P100 peak of the visual evoked potential. In the literature (see Di Russo et al. 2002), this component is also known as P1 or C2 (Jeffreys and Axford 1972a; Jeffreys and Axford 1972b). This component has been observed to have an onset latency between 65–80 ms and a peak latency between 100–130 ms; among our volunteers, we observed a peak latency of 108± 20 ms. Source localization was performed using the PCA-corrected reciprocal lead-field matrix *L* with inverse operator (cf. Knösche and Haueisen 2022):

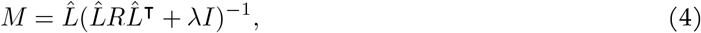

where 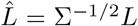 is the lead-field matrix whitened by the noise covariance matrix Σ, R is the source covariance matrix, and *λ* is a regularization parameter. In our case, the source-covariance matrix was chosen to be the diagonal matrix with

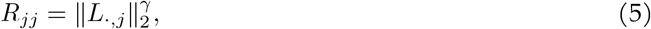

where *L*_*·,j*_ is the *j*-th column of the lead-field matrix, and *γ* is a depth-correction parameter. Localized source strengths are then obtained by applying the inverse operator to the whitened EEG signal Σ^-1*/*2^*v*. Source estimates were subsequently normalized using dSPM (Dale et al. 2000). The reader can find more details on our source reconstruction methodology in the papers (Nuñez Ponasso et al. 2025) and (Drumm et al. 2025).

## 3 Results

### 3.1 Source Estimation of Visual Evoked Potentials

The iPAF for each of the volunteers ranged between 8.75 –12 Hz. The P100 peak latencies we observed belonged to the range 84–155 ms; early and late phases had been previously observed in Di Russo et al. 2002 for pattern reversal stimuli.

In Figures 3 and 4, we highlight the dSPM P100 peak source reconstruction results for volunteers P01 and P20. The source reconstruction figures for the complete cohort can be found in Appendix B. In all cases, we displayed the reconstructed source strength density over the white matter surface and inflated white matter surface (the inflated white matter surfaces were created from the headreco meshes using our in-house implementation of the algorithm described in Fischl et al. 1999 with minor modifications). The reconstructed strengths were displayed in absolute value and normalized to the interval [0, 1]. The depth weighing parameter was chosen to be *γ* = 1.4 for all volunteers. Data for all 12 volunteers except for P10, P13, P14, and P16 was reconstructed with regularization parameter *λ* = 0.8; for these other volunteers we increased regularization parameter values according to the inverse operator’s condition number.

**Figure 3:**
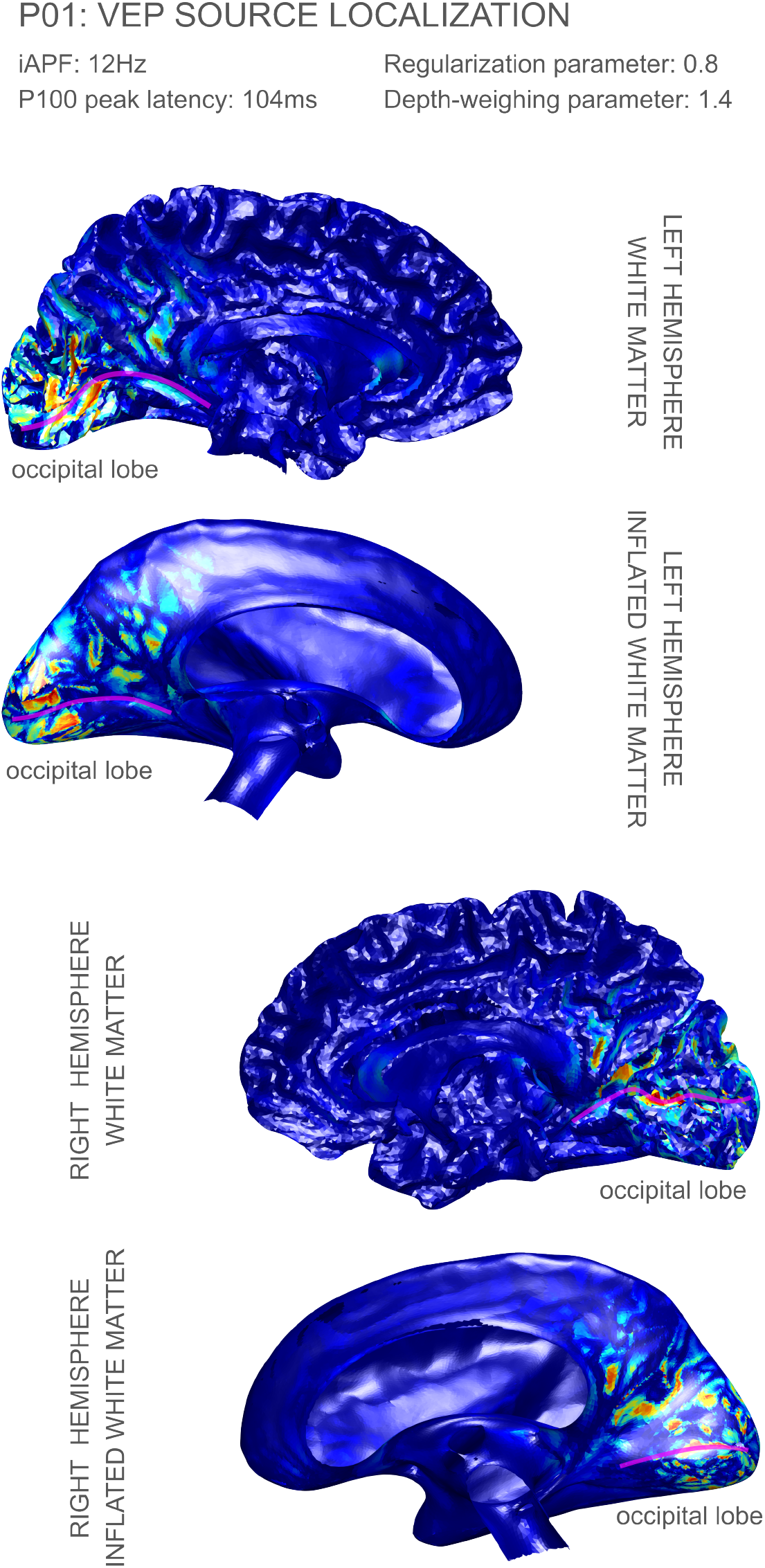
Source localization results for volunteer P01: the calcarine sulcus has been tagged in the WM and inflated WM surfaces with a magenta line. The corresponding P100 peaks of activation appeared at latency 104 ms, respectively. The regularization and depth parameters were λ = 0.8 and *γ* = 1.4, respectively.

**Figure 4:**
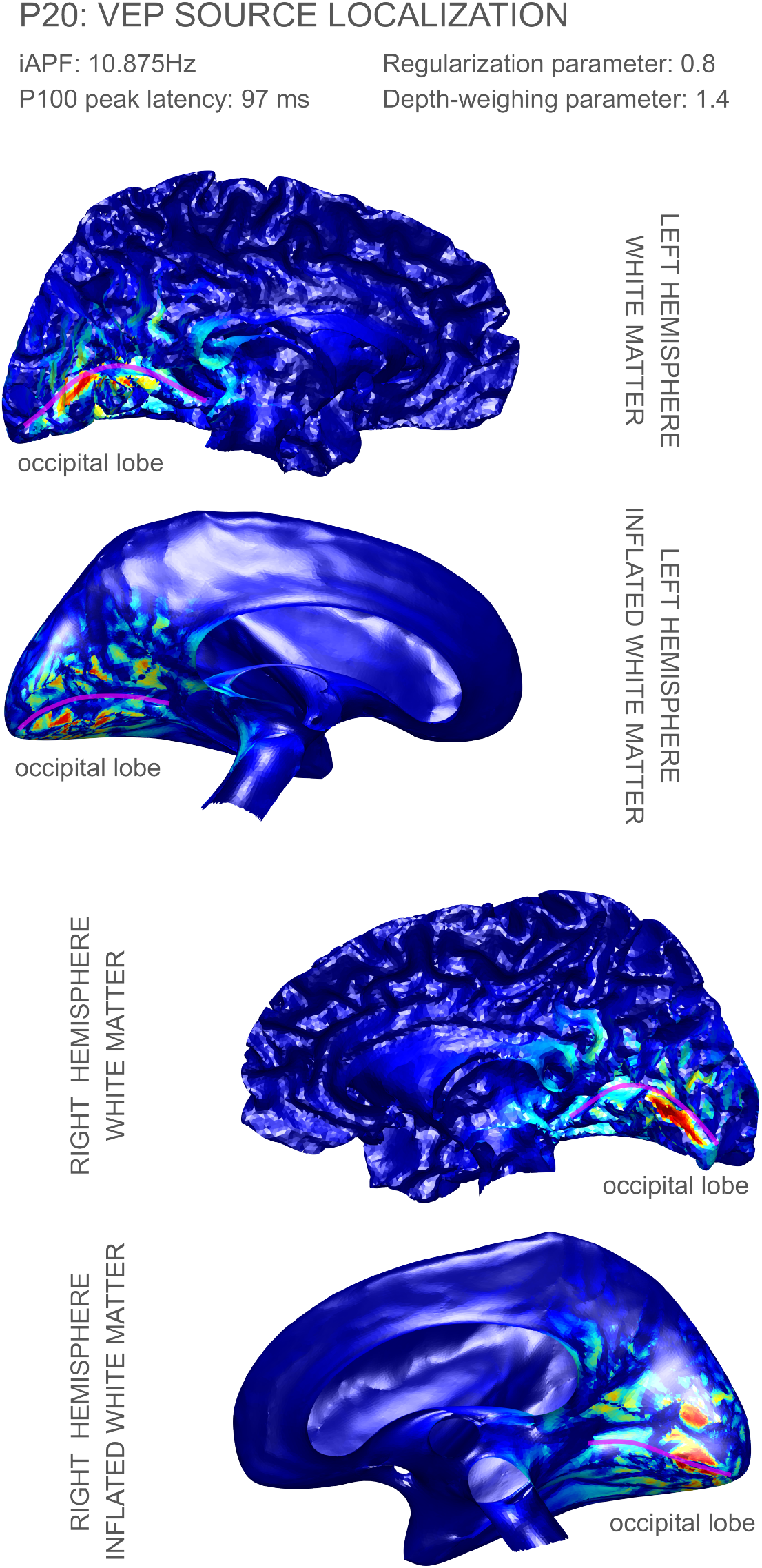
Source localization results for volunteer P20: the calcarine sulcus has been tagged in the WM and inflated WM surfaces with a magenta line. The corresponding P100 peaks of activation appeared at latency 97 ms, respectively. The regularization and depth parameters were λ = 0.8 and *γ* = 1.4, respectively.

## 4 Discussion

All source localization figures (cf. Figures 3 and 4, and the figures in Appendix B) are displayed on each individual’s left and right white matter hemispheres. All reconstruction results showed either no activation or negligible activation on the lateral regions of the cortex which are not displayed.

Both the localization results in Figures 3 and 4 (participants P01 and P20, respectively) show focal activation along the calcarine fissure (Brodmann area 17), and some less focal activation along Brodmann area 18 in the cuneus, with this secondary area of activation more pronounced in volunteer P01. Any possible components in Brodmann area 19 would be very weak on these two volunteers.

Figures in Appendix B, show activation peaks on area 17 in a consistent manner, although sometimes not in a symmetric way (like we observed in P01 and P20). For example, volunteer P14 shows very focal activation on the posterior part of area 17 (V1) in the left hemisphere while no significant amounts of activation in the right hemisphere. Another example of asymmetry is volunteer P21: in this case we observed very focal activation on the anterior part of area 17 in the right hemisphere, while most of the activity in the left hemisphere appears in area 18.

Overall our results show strong evidence supporting components of the P100 peak in Brodmann areas 17 and 18, with less evidence for area 19. This is consistent with the results for the P1 peak of the checkerboard stimulus in some of the papers surveyed in Di Russo et al. 2002. There has been a lack of consensus on the cortical visual areas generating P1, which can be attributed to limitations in the resolution of EEG systems, and the employed reconstruction methods. It should be noted, however, that those earlier results concern a different type of visual stimulus.

The effect of EEG cap density on our methods are currently unexplored. An interesting direction of future research would be to apply our methodology to experimental data obtained with high-density EEG systems (Fiedler et al. 2022).

Although the reciprocal approach is computationally much more efficient than the direct approach, it comes with an important drawback: whenever the sensor setup is changed, the BEM-FMM computation needs to be performed again. This is indeed a common experimental practice, and the general user may need to create several lead-field matrices when applying our methodology.

The practicability of a BCI systems significantly depends on the accuracy of the classification tasks. Today, BCIs operating in sensor space are already yielding promising results. However, sensor space-based BCI systems perform worse the more complex the classification tasks are. For example, (Ding et al. 2025) reported an average classification accuracy of 80.56% for a 2-finger MI control, but only a 60.61% accuracy for a 3-finger MI control. Applying source reconstruction methods enable the user to get more detailed and deeper insight into brain functioning. Handiru et al. 2016 describe an increase in the classification accuracy of multi-direction hand movements of more than 10% using source space features compared to sensor space features. But a key aspect that makes the use of source space analysis in BCI systems more difficult is the require real-time, or at least online capability of the algorithms. The BEM-FMM method presented here provides this capability after the lead-field matrix is calculated once. The current version of our algorithms allows for computation of an accurate and well conditioned inverse operator in less than 1 sec. Further, once the inverse operator is computed, the calculation of source reconstruction features can be performed near instantaneously (less than 0.1 sec), even for models with over 250,000 dipoles. This enables the utilization of our methodology for the control of external devices or communication applications within BCI systems.

## 5 Conclusions

We obtained remarkably focal source localization results for the P100/P1 component of visual evoked potentials. These results were obtained with 7 high-resolution realistic tissue meshes and involved ∼250,000 dipolar sources. Activation estimates for all subjects included a component in the calcarine sulcus (Brodmann area 17), as well as activity in area 18. Improvements in EEG source localization accuracy can have important implications in the field of BCIs.

## Acknowledgments

G.N.P., D.A.D., A.W, G.N., and S.N.M. were supported by the NIBIB Grant 1R01EB035484, and the NIMH Grant 1R01MH130490. H.O. received funding from the Thuringian Ministry for Economic Affairs, Science and Digital Society within the “Learning Products” project (5575/10-3). B.M. was supported by the German Federal Ministry of Education and Research (BMBF) grant DryPole (01GQ2304B).

J.H. received funding from the German Federal Ministry of Education and Research (BMBF) grant DryPole (01GQ2304A) and the Free State of Thuringia (2018 IZN 004), co-financed by the European Union under the European Regional Development Fund (ERDF).

## Appendices

### A. Derivation of Helmholtz Reciprocity for Current Electrodes and EEG

We derive in detail the EEG reciprocity relation for current electrodes and point dipoles. This is following and expanding upon the exposition of Plonsey 1963.

#### Derivation of quasi-static boundary value problems in bioelectromaggnetism

In bioelectromagnetism, the use of the quasi-static approximation to Maxwell’s Equations is justified (Plonsey and Heppner 1967). In practice, this means that we may neglect all time derivatives.

Denote the volume conductor (human head) by Ω and its surface by *∈*Ω. We will consider two types of problems, one with source currents only in the interior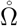 (case 1) of the head and another with source currents only on the surface *∂*Ω (case 2) of the head.

##### Case 1. Sources only in the interior of the head (EEG forward)

Let 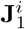be a primary, or impressed, current density which is zero. The total current **J**_1_ is split into two components

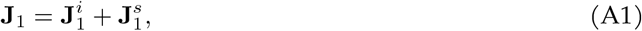

where 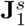is the secondary current, or volume current. In the case of EEG, the primary current can be approximated by a mathematical point-dipole in free space at a location **p** (de Munck et al. 1988):

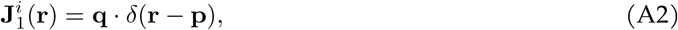

where **q** is the dipole moment. In particular, the primary field is everywhere continuous except at the singular point **p**. By Ohm’s Law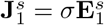, where *σ* can be regarded as a 3-tensor ℝ^3^ → ℝ^3^ × ℝ^3^, and 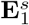 is the volume electric field, i.e. the electric field elicited by the deposited charge density in the presence of the primary current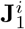. Namely, 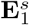is obtained from a charge density *ρ* via Coulomb’s Law, and as such this field is conservative and we can write 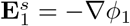for some scalar electric potential *ϕ*_1_. Applying the quasi-static approximation to the continuity equation yields:

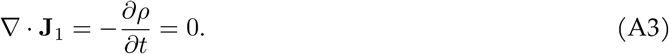

Combining Equations A1 and A3, we arrive at the fundamental equation:

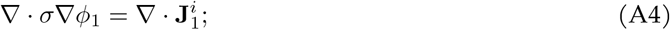

which must be satisfied everywhere in the interior of Ω.

A boundary condition can be also obtained from the continuity equation. First, note that we may approximate the conductivity of air as 0, so the skin surface *S* is a discontinuity surface for *σ*. Consider a Gaussian pillbox volume *V* centered at a point **r** ∈*S*, then the integral form of the continuity equation implies:

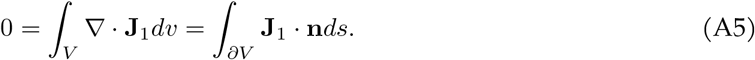

Letting the thickness of the pillbox approach 0, we find

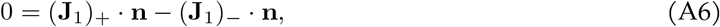

where (**J**_1_) *±*denotes the limiting values of **J**_1_ just outside and just inside of the skin surface *S*. In other words, the normal component of the total current is continuous across the surface *∂*Ω. We assumed that there are no impressed currents on the surface *∂*Ω, and there are no volume current outside of Ω, therefore (**J**_1_)_+_ = 0 and we arrive at the boundary condition

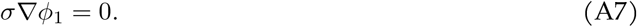

Putting everything together, we arrive at the following boundary value problem (which in particular governs EEG forward solutions):

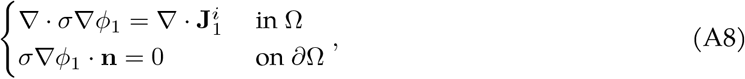

where *ϕ*_1_ is the scalar potential of the conservative field induced by the charge density in Ω in presence of the impressed current 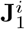.

##### Case 2. Impressed sources only on the head surface (TES forward)

We can replicate the same arguments as above, now with an impressed current 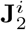which is zero every-where in the interior of Ω and non-zero only at some parts of the skin surface *∂*Ω. For example, in the case of TES we may take the following impressed current density

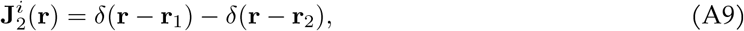

where **r**_1_ and **r**_2_ are the centers of two voltage electrodes on the skin surface.

The only difference is that now, 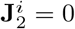in Ω so the differential equation in the interior is

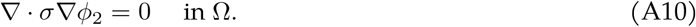

And on the surface we have that the total current just outside is precisely given by the impressed current 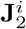(the current impressed by the electrodes on the skin surface). Hence, we obtain the boundary condition

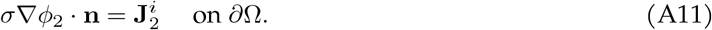

Which put together reads as the following boundary value problem:

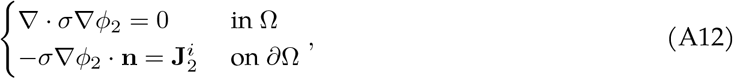

where *ϕ*_2_ is the scalar potential of the conservative field induced by the charge density in Ω in presence of the impressed current 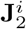. Here, we are assuming that the normals of *∂*Ω are directed inwards (hence the negative sign appearing).

##### Green’s Theorem

We have the following identities for the “cross-terms” of the two potentials *ϕ*_1_ and *ϕ*_2_ described above:

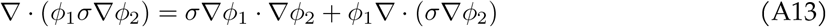

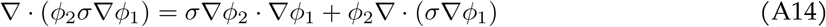

Applying Gauss’s Divergence Theorem, and subtracting we obtain

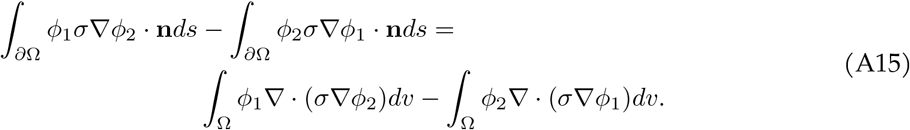

Now, applying the conditions in Equations A8 and A12 to *ϕ*_1_ and *ϕ*_2_ respectively, we find

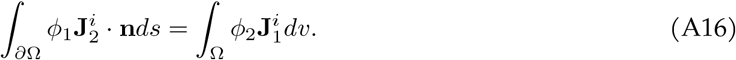

This is the continuous version of Green’s Theorem.

##### EEG-TES reciprocity

We now let

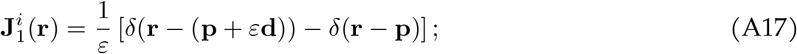

here *ε* is a dimensionless scaling constant. This is the impressed current density of a unit-strength finite length current dipole with displacement direction **d** (a unit vector) and length *ε >* 0.

On the other hand, we let

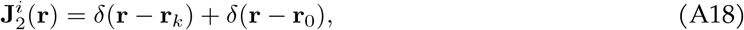

where **r**_*k*_ is the center of the *k*-th electrode of an EEG sensor sytem, and **r**_0_ is the reference electrode. Substituting into Equation A16, we find

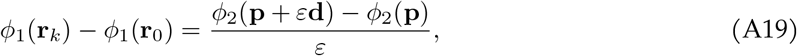

which holds true for all *ε >* 0. Note that this is an identity only involving potential differences, so it is consistent in terms of units. Now, using the definition of the derivative, we have

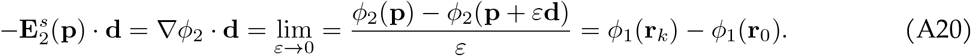

The leftmost term is in units [V m^-1^ m]=[V], so this equation is also consistent in terms of units. We arrive at the fundamental Helmholtz’s reciprocity relating EEG forward solutions with TES forward solutions:

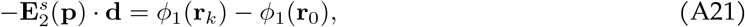

where 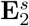is the volume field elicited by TES at cathode position **r**_*k*_ and anode position **r**_0_, and *ϕ*_1_ is the potential of the volume field elicited by EEG with a point dipole of unit direction **d** located at position **p**.

**Figure A1.**
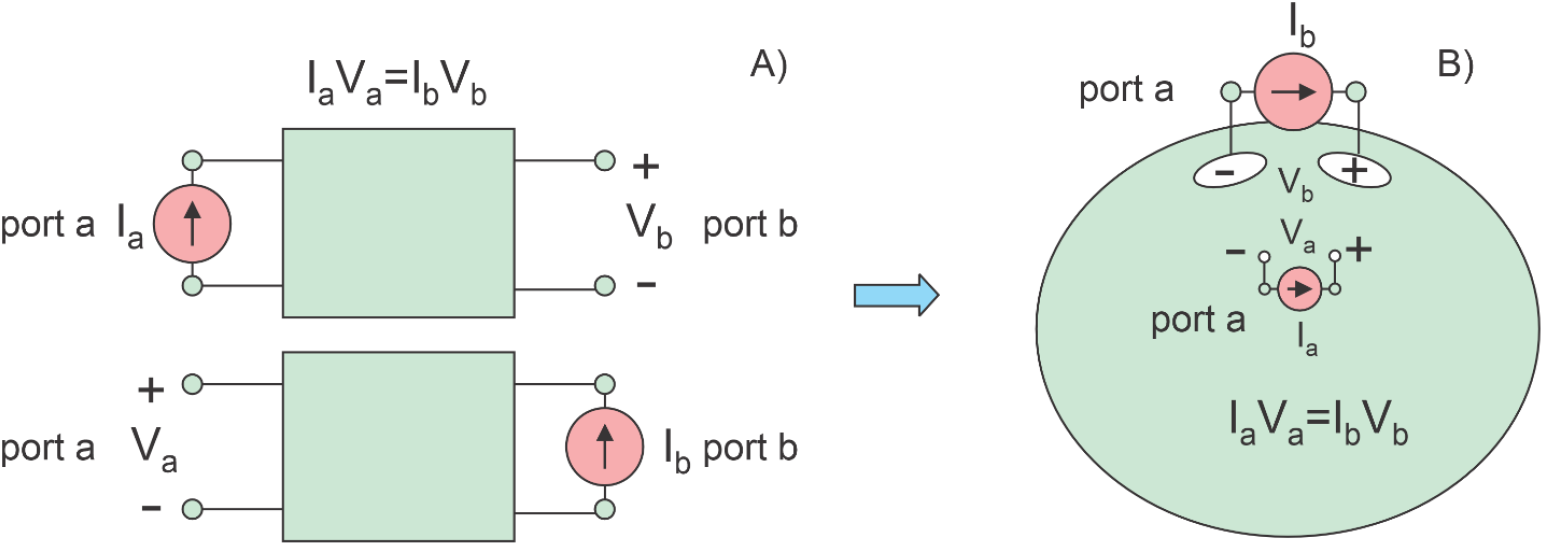
A. Circuital reciprocity theorem. B. Its application to a distributed circuit – a volume conductor.

#### A.1 EEG Reciprocity from Circuit Reciprocity

The quasistatic reciprocity (duality) principle or theorem (Helmholtz 1853; Rush and Driscoll 1969) makes it possible to interrelate the fields within the head generated by surface electrodes with the fields of the forward EEG problem with embedded dipolar point current sources (Vallaghé et al. 2008; Wagner et al. 2016). Also, the corresponding powerful analytical EEG solutions — cf. (Zhang 1995; Mosher et al. 1999) — could be used to test the validity of the fields injected by surface electrodes. In the relevant EEG studies (Wagner et al. 2016; Vanrumste et al. 2001; Laarne et al. 2000; Finke et al. 2013; Fletcher et al. 1995; Shirvany et al. 2013) the reciprocal solution was primarily used to more easily obtain multiple forward solutions with the elementary dipoles. Then, these forward solutions were employed in a standard framework of the EEG inverse analysis. We apply a somewhat different approach. It effectively bypasses the forward EEG solutions and utilizes the fields generated by surface electrodes for the solution of the inverse EEG problem over the entire cortical surface.

##### A.1.1 One electrode pair and one cortical dipole

Among other methods, a straightforward way to derive the necessary relation is to directly apply the circuital reciprocity theorem (Lorrain et al. 1987; Guillemin 1953) for a linear passive circuit. The circuit is connected to a current source and a voltmeter as in Fig. 1A. In this case, interchanging the ideal current source and the voltmeter does not alter the ratio V/I so that *V*_*a*_*I*_*a*_ = *V*_*b*_*I*_*b*_ (Lorrain et al. 1987; Guillemin 1953).

Nothing changes if the linear circuit takes the form of a volume conductor with many distributed resistances (Fig. A1B). We set Port a (a dipole current source) somewhere inside the conductor while Port b (a pair of surface current electrodes) will be located at its surface. ·In terms of these notations, the equality *V*_*a*_*I*_*a*_ = *V*_*b*_*I*_*b*_ becomes

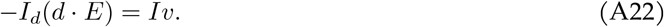

##### A.1.2 One electrode pair and multiple cortical dipoles

We keep the surface current source in Fig. A1B *the same* but try out different internal dipole sources at different locations *r*_*n*_, *n* = 1, …, *N*. In other words, we turn these cortical dipoles on, one by one, and obtain, from Eq. A22

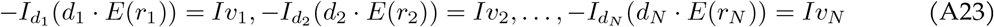

Adding up all Eqs. A23 yields

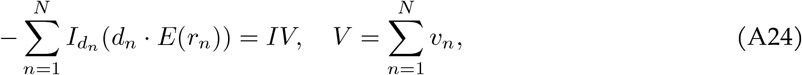

where *V* is the total surface electrode voltage generated by the dipole layer when all dipoles are turned on. The continuous (integral) version of Eq. A24 has the form

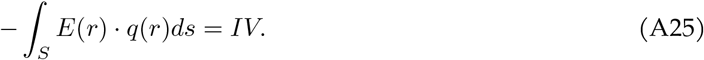

Here, *q*(*r*) (*A* ·*m/m*^2^) is the current dipole moment per unit cross sectional area of a continuous dipole layer with surface *S, E*(*r*) (V/m) is the continuous field of the surface current source across the dipole layer, *I* (A) is the strength of the surface current source and *V* (V) is the voltage between the two surface electrodes generated by the dipole layer.

The reciprocity principle is only applicable to a single surface electrode pair at a time. All other electrode pairs, i.e., all other surface current sources must be turned off.

### B Additional Figures

**Figure B1:**
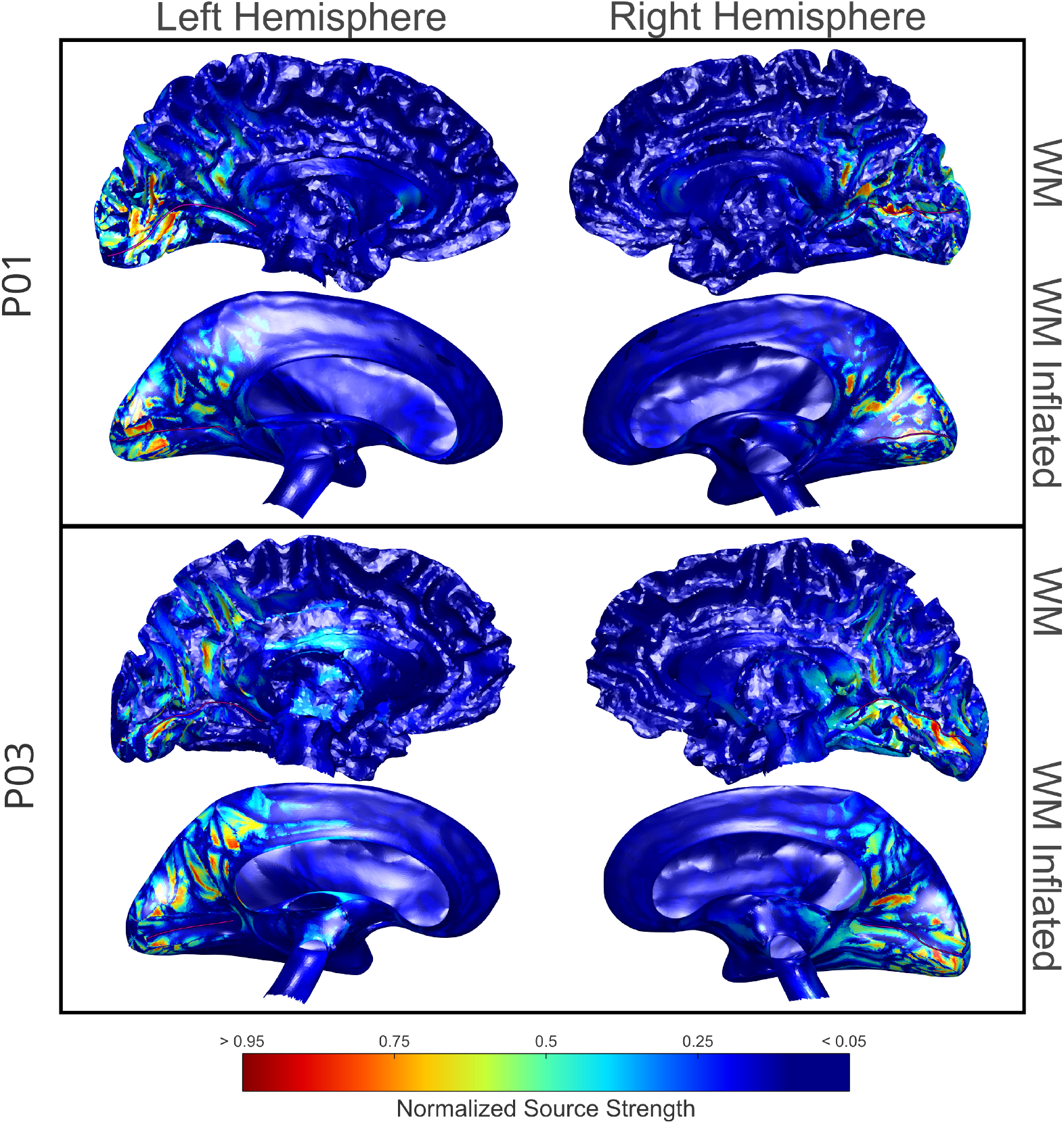
Source localization results for volunteers P01 and P03: the calcarine sulcus has been tagged in the WM and inflated WM surfaces with a magenta line. The corresponding P100 peaks of activation appeared in latencies 104 and 111 ms, respectively. The regularization and depth parameters were *λ* = 0.8 and *γ* = 1.4 for both volunteers.

**Figure B2:**
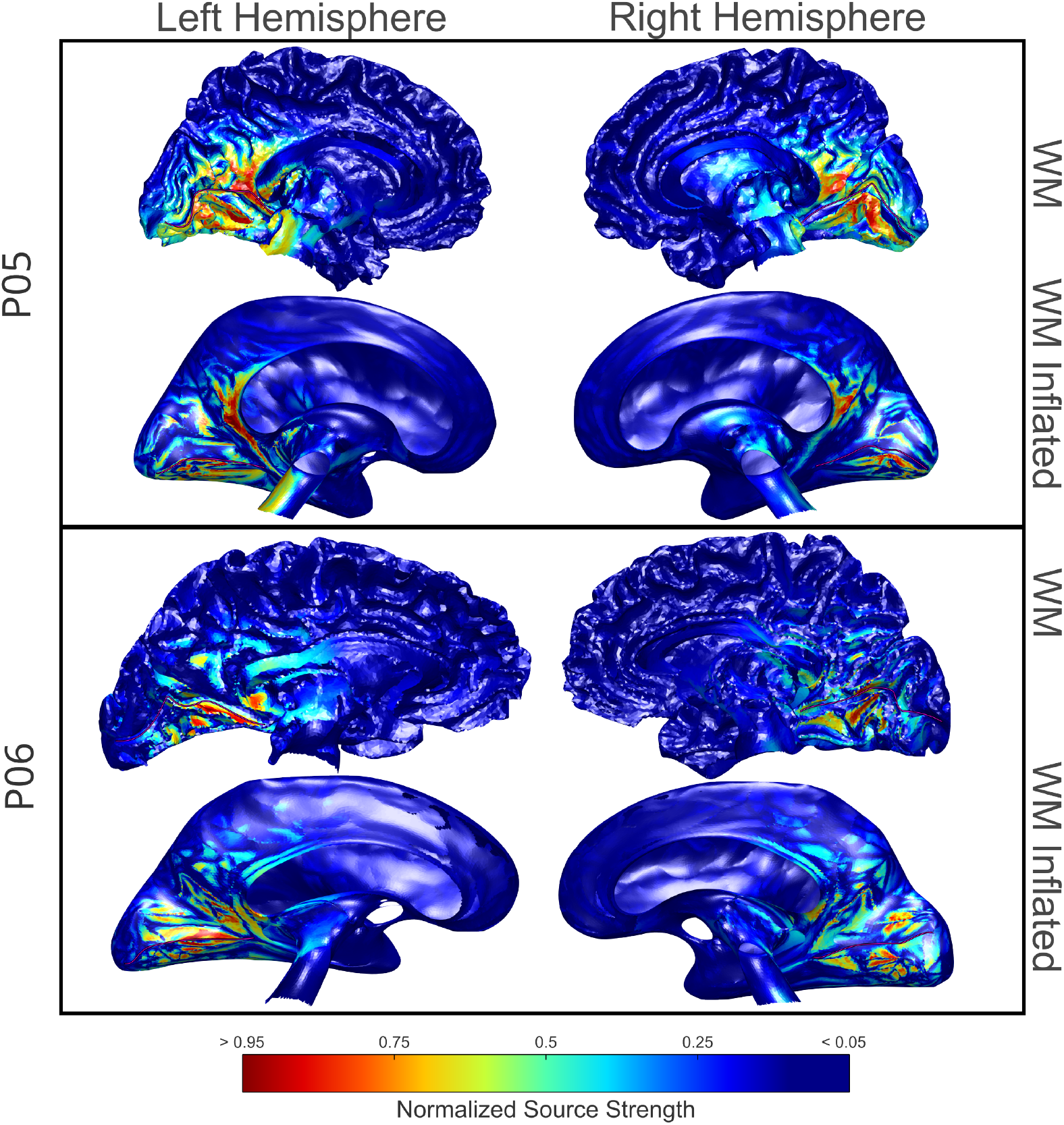
Source localization results for volunteers P05 and P06: the calcarine sulcus has been tagged in the WM and inflated WM surfaces with a magenta line. The corresponding P100 peaks of activation appeared in latencies 99 and 133 ms, respectively. The regularization and depth parameters were *λ* = 0.8 and *γ* = 1.4 for both volunteers.

**Figure B3:**
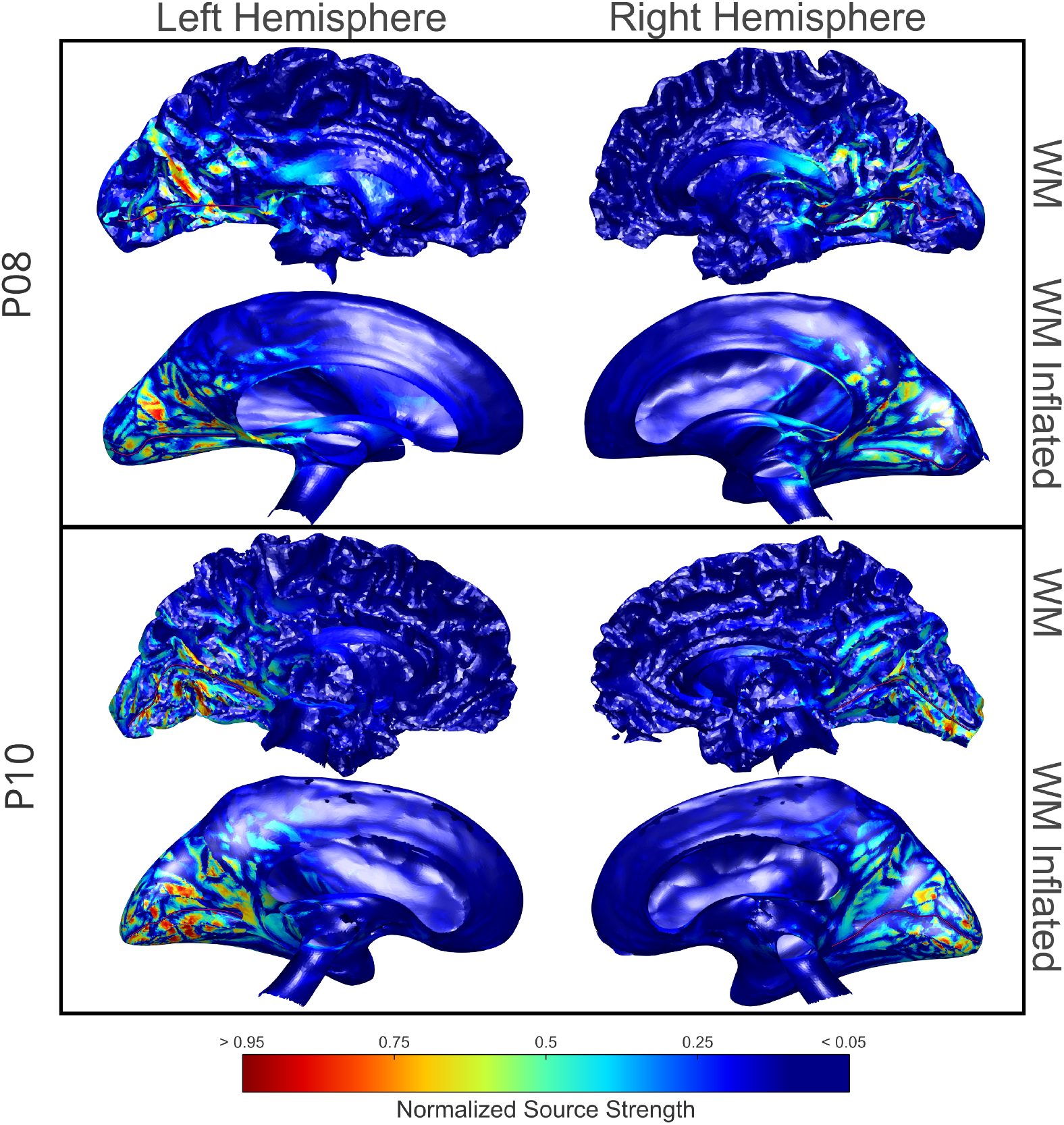
Source localization results for volunteers P08 and P10: the calcarine sulcus has been tagged in the WM and inflated WM surfaces with a magenta line. The corresponding P100 peaks of activation appeared in latencies 103 and 84 ms, respectively. The regularization parameters were *λ* = 0.8 and *λ* = 2 for each volunteer, respectively. The depth parameter *γ* = 1.4 was used for both volunteers.

**Figure B4:**
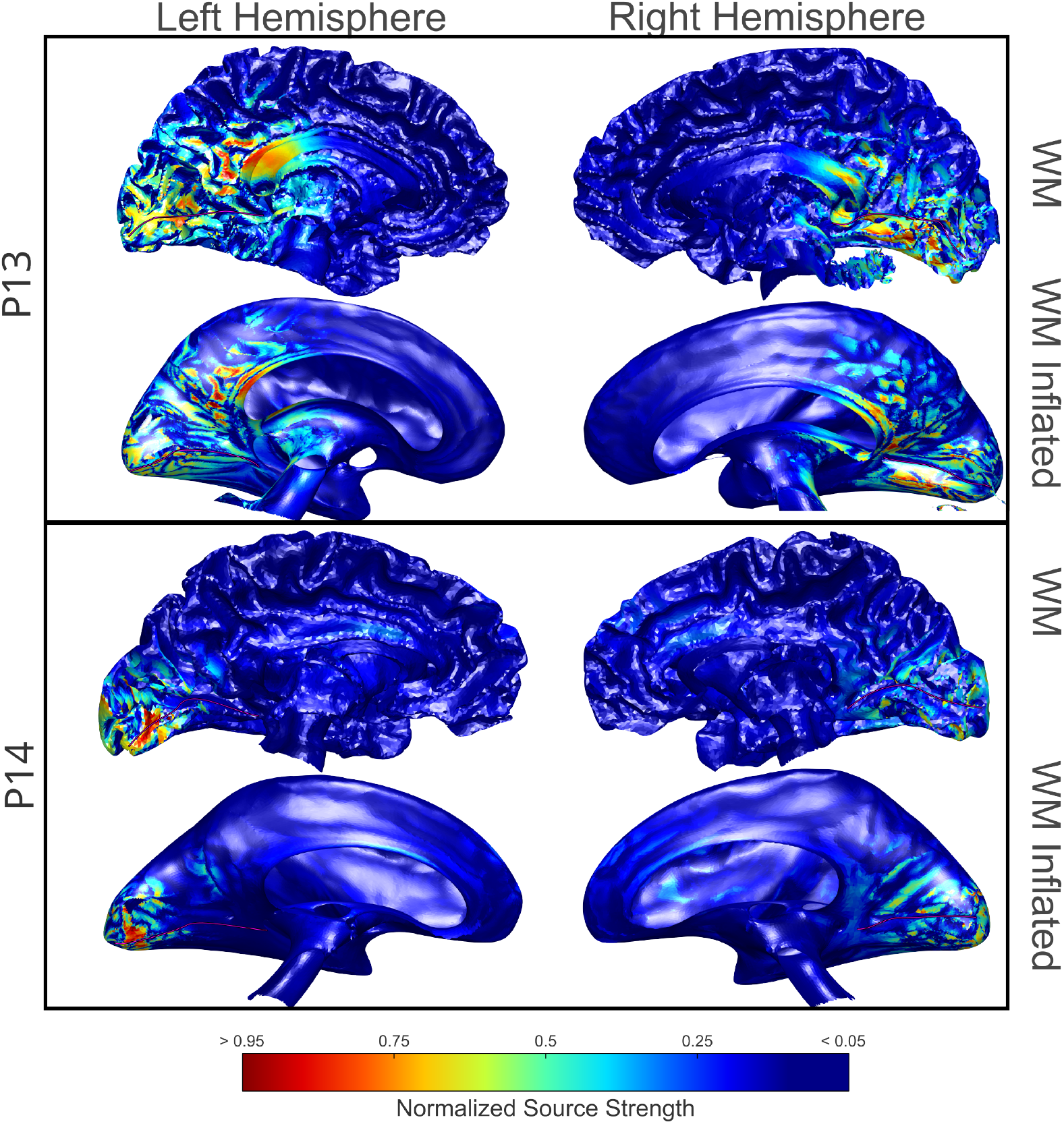
Source localization results for volunteers P13 and P14: the calcarine sulcus has been tagged in the WM and inflated WM surfaces with a magenta line. The corresponding P100 peaks of activation appeared in latencies 90 and 85 ms, respectively. The regularization parameters were *λ* = 1.5 and *λ* = 0.8 for each volunteer, respectively. The depth parameter *γ* = 1.4 was used for both volunteers.

**Figure B5:**
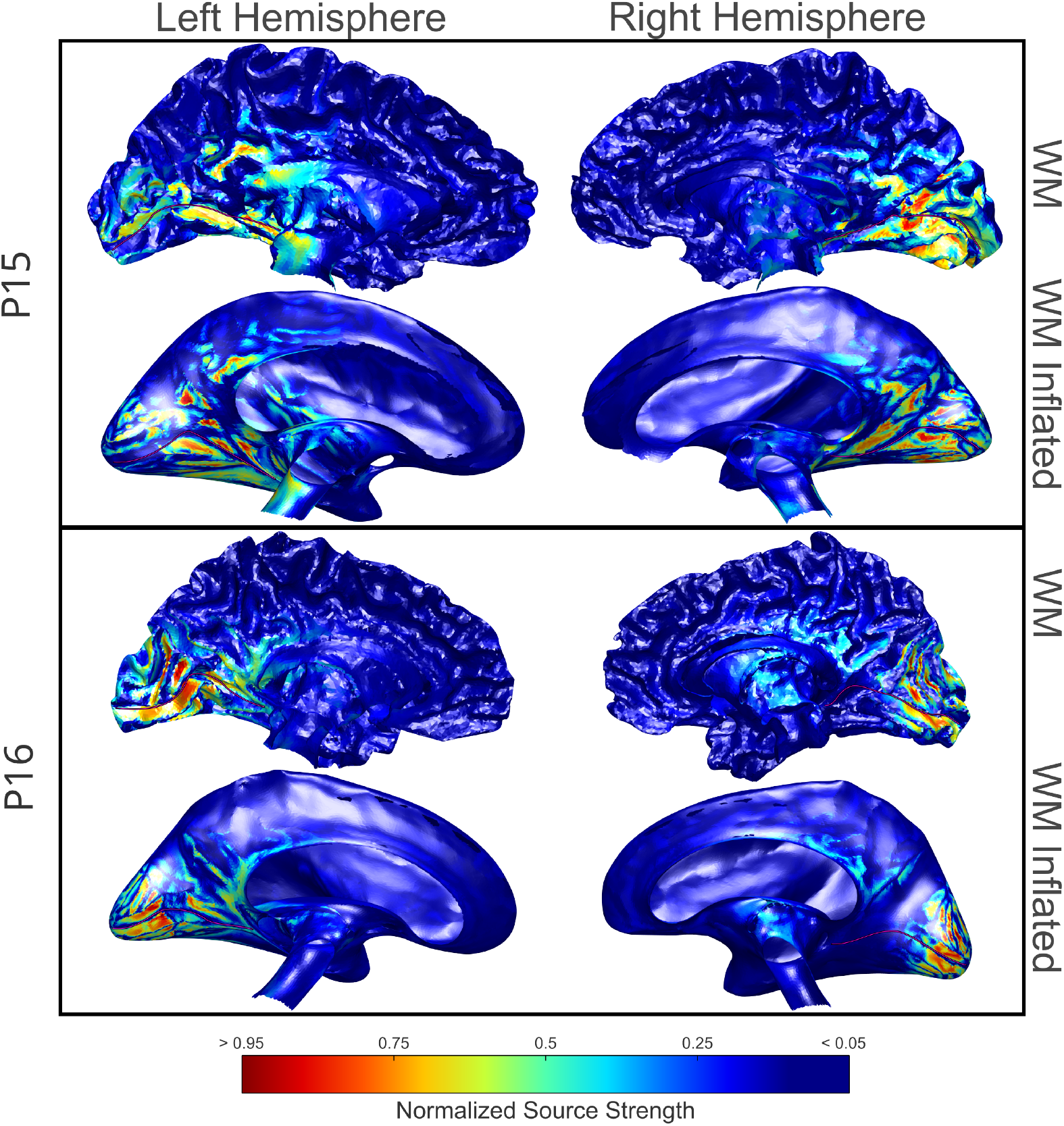
Source localization results for volunteers P15 and P16: the calcarine sulcus has been tagged in the WM and inflated WM surfaces with a magenta line. The corresponding P100 peaks of activation appeared in latencies 117 and 122 ms, respectively. The regularization parameters were *λ* = 0.8 and *γ* = 1.4 for both volunteers.

**Figure B6:**
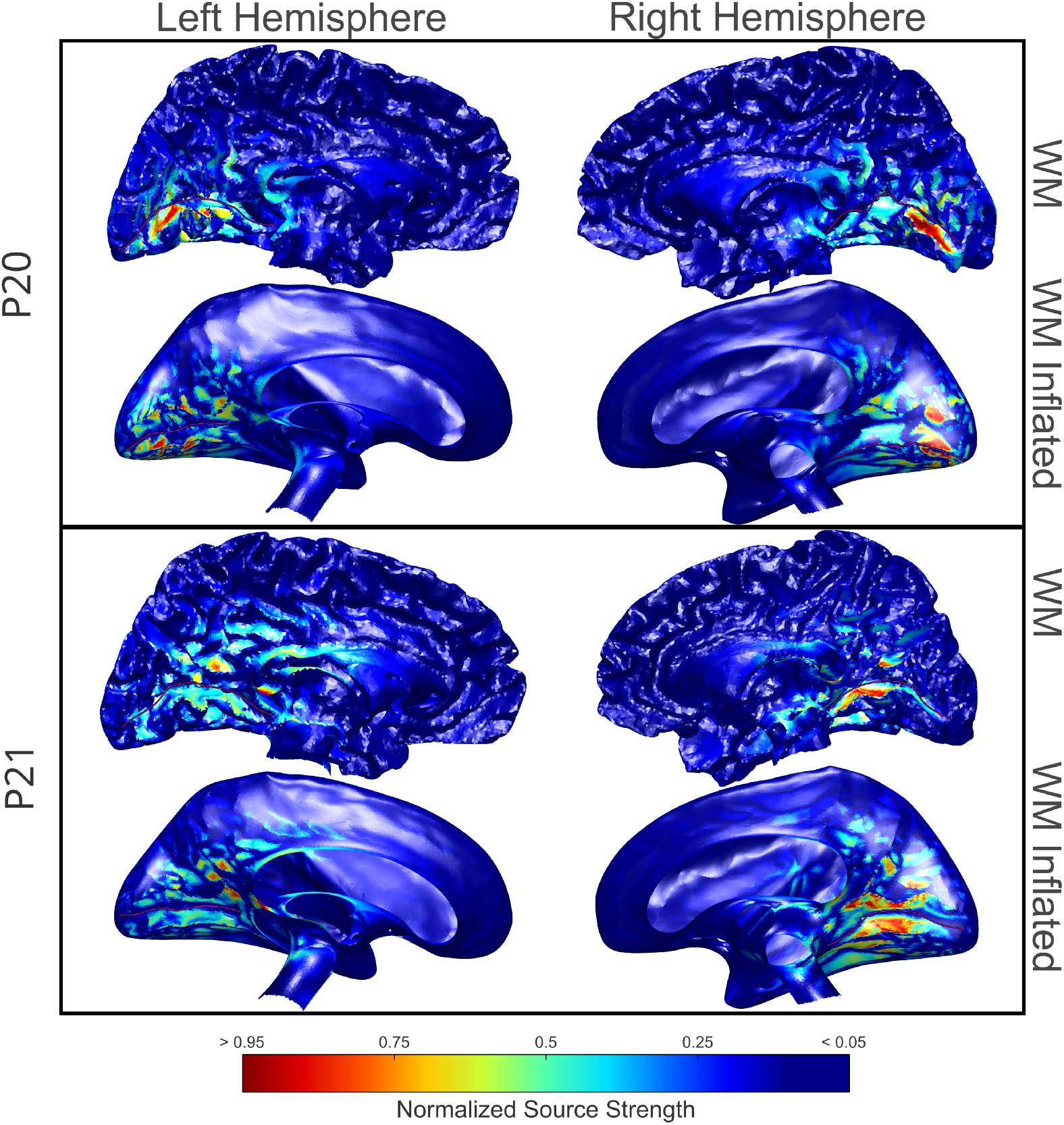
Source localization results for volunteers P20 and P21: the calcarine sulcus has been tagged in the WM and inflated WM surfaces with a magenta line. The corresponding P100 peaks of activation appeared in latencies 97 and 112 ms, respectively. The regularization parameters were *λ* = 0.8 and *λ* = 1.5 for each volunteer, respectively. The depth parameter *γ* = 1.4 was used for both volunteers.

## References

Ai, Jikun et al. (2023). “BCI Control of a Robotic Arm Based on SSVEP With Moving Stimuli for Reach and Grasp Tasks”. In: IEEE Journal of Biomedical and Health Informatics 27.8, pp. 3818–3829. doi: 10.1109/JBHI.2023.3277612.

Barnard, A.C. L. et al. (1967). “The Application of Electromagnetic Theory to Electrocardiology: I. Derivation of the Integral Equations”. In: Biophysical Journal 7.5, pp. 443–462. issn: 0006-3495. doi: 10.1016/S0006-3495(67)86598-6.

Chota, Samson et al. (2024). “Steady-state Visual Evoked Potentials Reveal Dynamic (Re)allocation of Spatial Attention during Maintenance and Utilization of Visual Working Memory”. In: Journal of Cognitive Neuroscience 36.5, pp. 800–814. issn: 0898-929X. doi: 10.1162/jocn_a_02107. URL: https://doi.org/10.1162/jocn%5C_a%5C_02107.

Compston, Alastair (2009). “The Berger rhythm: potential changes from the occipital lobes in man, by E.D. Adrian and B.H.C. Matthews (From the Physiological Laboratory, Cambridge). Brain 1934: 57; 355–385.” In: Brain 133.1, pp. 3–6. issn: 0006-8950. doi: 10.1093/brain/awp324. eprint: https://academic.oup.com/brain/article-pdf/133/1/3/894742/awp324.pdf. url: https://doi.org/10.1093/brain/awp324.

Dale, Anders M. et al. (2000). “Dynamic Statistical Parametric Mapping: Combining fMRI and MEG for High-Resolution Imaging of Cortical Activity”. In: Neuron 26.1, pp. 55–67. issn: 0896-6273. doi: 10.1016/S0896-6273(00)81138-1.

de Munck, J.C. et al. (1988). “Mathematical dipoles are adequate to describe realistic generators of human brain activity”. In: IEEE Trans. Biomed. Eng. 35.11, pp. 960–966. doi: 10.1109/10.8677.

Di Russo, Francesco et al. (2002). “Cortical sources of the early components of the visual evoked potential”. In: Human Brain Mapping 15.2, pp. 95–111. doi: 10.1002/hbm.10010.

Ding, Yidan et al. (2025). “EEG-based brain-computer interface enables real-time robotic hand control at individual finger level.” In: Nature Communications 16.1. doi: 10.1038/s41467-025-61064-x.

Drumm, Derek A. et al. (2025). “Improved Source Localization of Auditory Evoked Fields using Reciprocal BEM-FMM”. In: bioRxiv. doi: 10.1101/2025.05.09.653081. URL: https://www.biorxiv.org/content/early/2025/05/14/2025.05.09.653081.

Fiedler, Patrique et al. (2022). “A high-density 256-channel cap for dry electroencephalography”. In Human Brain Mapping 43.4, pp. 1295–1308. doi: 10.1002/hbm.25721.

Finke, Stefan et al. (2013). “Conventional and Reciprocal Approaches to the Inverse Dipole Localization Problem for N20–P20 Somatosensory Evoked Potentials”. In: Brain Topography 26.1, pp. 24–34. issn: 1573-6792. doi: 10.1007/s10548-012-0238-x.

Fischl, Bruce et al. (1999). “Cortical Surface-Based Analysis: II: Inflation, Flattening, and a Surface-Based Coordinate System”. In: NeuroImage 9.2, pp. 195–207. issn: 1053-8119. doi: 10.1006/nimg.1998.0396.

Fischl, Bruce et al. (2004). “Sequence-independent segmentation of magnetic resonance images”. In: NeuroImage 23. Mathematics in Brain Imaging, S69–S84. issn: 1053-8119. doi: 10.1016/j.neuroimage.2004.07.016.

Fletcher, D.J. et al. (1995). “Improved method for computation of potentials in a realistic head shape model”. In: IEEE Transactions on Biomedical Engineering 42.11, pp. 1094–1104. doi: 10.1109/10.469376.

Frobenius, Georg (1912). “Ueber Matrizen aus nicht negativen Elementen”. In: Sitzungsberichte der Königlich Preussischen Akademie der Wissenschaften, pp. 456–477.

Gelernter, H.L. and J.C. Swihart (1964). “A Mathematical-Physical Model of the Genesis of the Electro-cardiogram”. In: Biophys. J. 4.4, pp. 285–301. issn: 0006-3495. doi: 10.1016/S0006-3495(64)86783-7.

Geselowitz, D. (1970). “On the magnetic field generated outside an inhomogeneous volume conductor by internal current sources”. In: IEEE Transactions on Magnetics 6.2, pp. 346–347. doi: 10.1109/TMAG.1970.1066765.

Geselowitz, David B. (1967). “On Bioelectric Potentials in an Inhomogeneous Volume Conductor”. In: Biophys. J. 7.1, pp. 1–11. doi: 10.1016/S0006-3495(67)86571-8.

Gomez, Luis J. et al. (2020). “Conditions for numerically accurate TMS electric field simulation”. In: Brain Stimulation 13.1, pp. 157–166. issn: 1935-861X. doi: 10.1016/j.brs.2019.09.015.

Gramfort, Alexandre et al. (2010). “OpenMEEG: opensource software for quasistatic bioelectromagnetics”. In: BioMedical Engineering OnLine 9.1, p. 45. issn: 1475-925X. doi: 10.1186/1475-925X-9-45.

Gramfort, Alexandre et al. (2013). “MEG and EEG Data Analysis with MNE-Python”. In: Front. Neurosci. 7.267, pp. 1–13. doi: 10.3389/fnins.2013.00267.

Gramfort, Alexandre et al. (2014). “MNE Software for Processing MEG and EEG Data”. In: NeuroImage 86, pp. 446–460. doi: 10.1016/j.neuroimage.2013.10.027.

Greengard, L and V Rokhlin (1987). “A fast algorithm for particle simulations”. In: J. Comput. Phys. 73.2, pp. 325–348. issn: 0021-9991. doi: 10.1016/0021-9991(87)90140-9.

Guillemin, Ernst A. (1953). Introductory Circuit Theory. Section 6: The reciprocity theorem. New York: John Wiley & Sons. Chap. 3, pp. 148–153. ISBN: 0471330663.

Hämäläinen, M. S. and R. J. Ilmoniemi (1994). “Interpreting magnetic fields of the brain: minimum norm estimates”. In: Med. Biol. Eng. Comput. 32.1, pp. 35–42. issn: 1741-0444. doi: 10.1007/BF02512476.

Hämäläinen, Matti et al. (1993). “Magnetoencephalography—theory, instrumentation, and applications to noninvasive studies of the working human brain”. In: Rev. Mod. Phys. 65 (2), pp. 413–497. doi: 10.1103/RevModPhys.65.413.

Handiru, Vikram Shenoy et al. (2016). “Multi-direction hand movement classification using EEG-based source space analysis”. In: pp. 4551–4554. doi: 10.1109/EMBC.2016.7591740.

Helmholtz, H. (1853). “Ueber einige Gesetze der Vertheilung elektrischer Ströme in körperlichen Leitern, mit Anwendung auf die thierisch-elektrischen Versuche (Schluss.)” In: Ann. der Phys. (und Chemie) 165.7, pp. 353–377. doi: 10.1002/andp.18531650702.

Hestenes, M.R. and E. Stiefel (1952). “Methods of Conjugate Gradients for Solving Linear Systems”. In: J. Res. Natl. Bur. Stand. 49, pp. 409–435. doi: http://dx.doi.org/10.6028/jres.049.044.

The Flatiron Institute (2012). FMM3D: Flatiron Institute Fast Multipole Libraries. url: https://github.com/flatironinstitute/FMM3D.

IT’IS Foundation (2024). Automated Head Segmentation in Sim4Life V8.0.1: A New Reference. url: https://itis.swiss/s/news-events/news/virtual-population/ixi-heads.

Jeffreys, D. A. and J. G. Axford (1972a). “Source locations of pattern-specific components of human visual evoked potentials. I. Component of striate cortical origin”. In: Experimental Brain Research 16.1, pp. 1–21. issn: 1432-1106. doi: 10.1007/BF00233371.

Jeffreys, D. A. and J. G. Axford (1972b). “Source locations of pattern-specific components of human visual evoked potentials. II. Component of extrastriate cortical origin”. In: Experimental Brain Research 16.1, pp. 22–40. issn: 1432-1106. doi: 10.1007/BF00233372.

Knösche Thomas R. and Jens Haueisen (2022). EEG/MEG Source Reconstruction. Textbook for Electro-and Magnetoencephalography. Springer Cham. ISBN: 978-3-030-74916-3. doi: 10.1007/978-3-030-74918-7.

Kybic, J. et al. (2005). “A common formalism for the Integral formulations of the forward EEG problem”. In: IEEE Trans. Med. Imaging 24.1, pp. 12–28. doi: 10.1109/TMI.2004.837363.

Laarne, Päivi et al. (2000). “Accuracy of Two Dipolar Inverse Algorithms Applying Reciprocity for Forward Calculation”. In: Computers and Biomedical Research 33.3, pp. 172–185. issn: 0010-4809. doi: 10.1006/cbmr.1999.1538.

Li, Minglun et al. (2021). “Brain–Computer Interface Speller Based on Steady-State Visual Evoked Potential: A Review Focusing on the Stimulus Paradigm and Performance”. In: Brain Sciences 11.4. issn: 2076-3425. doi: 10.3390/brainsci11040450. url: https://www.mdpi.com/2076-3425/11/4/450.

Li, Yuanqing et al. (2013). “A Hybrid BCI System Combining P300 and SSVEP and Its Application to Wheelchair Control”. In: IEEE Transactions on Biomedical Engineering 60.11, pp. 3156–3166. doi: 10.1109/TBME.2013.2270283.

Lin, Fa-Hsuan et al. (2006). “Distributed current estimates using cortical orientation constraints”. In: Human Brain Mapping 27.1, pp. 1–13. doi: 10.1002/hbm.20155. eprint: https://onlinelibrary.wiley.com/doi/pdf/10.1002/hbm.20155.

Lorrain, Paul et al. (1987). Electromagnetic Fields and Waves Including Electric Circuits. 3rd. W. H. Freeman & Co. Chap. 8, pp. 157–161. isbn: 0716718235.

Makarov, Sergey N et al. (2020). “A software toolkit for TMS electric-field modeling with boundary element fast multipole method: an efficient MATLAB implementation”. In: J. Neural Eng. 17.4, p. 046023. doi: 10.1088/1741-2552/ab85b3.

Makarov, Sergey N et al. (2021a). “Boundary element fast multipole method for modeling electrical brain stimulation with voltage and current electrodes”. In: J. Neural Eng. 18.4, p. 0460d4. doi: 10.1088/1741-2552/ac17d7.

Makarov, Sergey N. et al. (2018). “A Quasi-Static Boundary Element Approach With Fast Multipole Acceleration for High-Resolution Bioelectromagnetic Models”. In: IEEE Trans. Biomed. Eng. 65.12, pp. 2675–2683. doi: 10.1109/TBME.2018.2813261.

Makarov, Sergey N. et al. (2021b). “Boundary Element Fast Multipole Method for Enhanced Modeling of Neurophysiological Recordings”. In: IEEE Trans. Biomed. Eng. 68.1, pp. 308–318. doi: 10.1109/TBME.2020.2999271.

Mosher, J.C. et al. (1999). “EEG and MEG: forward solutions for inverse methods”. In: IEEE Transactions on Biomedical Engineering 46.3, pp. 245–259. doi: 10.1109/10.748978.

Nielsen, Jesper D. et al. (2018). “Automatic skull segmentation from MR images for realistic volume conductor models of the head: Assessment of the state-of-the-art”. In: NeuroImage 174, pp. 587–598. issn: 1053-8119. doi: 10.1016/j.neuroimage.2018.03.001. URL: https://www.sciencedirect.com/science/article/pii/S1053811918301800.

Norcia, Anthony M. et al. (2015). “The steady-state visual evoked potential in vision research: A review”. In: Journal of Vision 15.6, pp. 4–4. issn: 1534-7362. doi: 10.1167/15.6.4. URL: https://doi.org/10.1167/15.6.4.

Nunez, Paul L. (1981). Electric Fields of the Brain: the Neurophysics of EEG. New York: Oxford University Press.

Nuñez Ponasso, Guillermo (2024). “A survey on integral equations for bioelectric modeling”. In: Phys. Med. Biol. 69.17, 17TR02. doi: 10.1088/1361-6560/ad66a9.

Nuñez Ponasso, Guillermo et al. (2025). “High-Definition MEG Source Estimation using the Reciprocal Boundary Element Fast Multipole Method”. In: bioRxiv. doi: 10.1101/2025.03.21.644601. URL: https://www.biorxiv.org/content/early/2025/03/24/2025.03.21.644601.

Oikonomou, Vangelis P. and Ioannis Kompatsiaris (2020). “A Novel Bayesian Approach for EEG Source Localization”. In: Computational Intelligence and Neuroscience 2020.1, p. 8837954. doi: 10.1155/2020/8837954. URL: https://onlinelibrary.wiley.com/doi/abs/10.1155/2020/8837954.

Oostenveld, Robert et al. (2011). “FieldTrip: Open Source Software for Advanced Analysis of MEG, EEG, and Invasive Electrophysiological Data”. In: Comput. Intell. Neurosci. 2011.1, p. 156869. doi: 10.1155/2011/156869.

Perron, Oskar (1907). “Zur Theorie der Matrices”. In: Mathematische Annalen 64.2, pp. 248–263. issn: 1432-1807. doi: 10.1007/BF01449896.

Plonsey, Robert (1963). “Reciprocity Applied to Volume Conductors and the ECG”. In: IEEE Transactions on Bio-medical Electronics 10.1, pp. 9–12. doi: 10.1109/TBMEL.1963.4322775.

Plonsey, Robert and Dennis B. Heppner (1967). “Considerations of quasi-stationarity in electrophysiological systems”. In: Bull. Math. Biophys. 29.4, pp. 657–664. issn: 1522-9602. doi: 10.1007/BF02476917.

Rush, Stanley and Daniel A. Driscoll (1969). “EEG Electrode Sensitivity-An Application of Reciprocity”. In: IEEE Transactions on Biomedical Engineering BME-16.1, pp. 15–22. doi: 10.1109/TBME.1969.4502598.

Saturnino, Guilherme B. et al. (2019). “SimNIBS 2.1: A Comprehensive Pipeline for Individualized Electric Field Modelling for Transcranial Brain Stimulation”. In: Brain and Human Body Modeling: Computational Human Modeling at EMBC 2018. Ed. by Sergey Makarov et al. Cham: Springer International Publishing, pp. 3–25. ISBN: 978-3-030-21293-3. doi: 10.1007/978-3-030-21293-3_1.

Savitzky, Abraham. and M. J. E. Golay (1964). “Smoothing and Differentiation of Data by Simplified Least Squares Procedures.” In: Anal. Chem. 36.8, pp. 1627–1639. doi: 10.1021/ac60214a047.

Schielke, Alexander and Bart Krekelberg (2022). “Steady state visual evoked potentials in schizophrenia: A review”. In: Frontiers in Neuroscience Volume 16 - 2022. issn: 1662-453X. doi: 10.3389/fnins.2022.988077. URL: https://www.frontiersin.org/journals/neuroscience/articles/10.3389/fnins.2022.988077.

Schrader, Sophie et al. (2021). “DUNEuro—A software toolbox for forward modeling in bioelectromagnetism”. In: PLOS ONE 16.6, pp. 1–21. doi: 10.1371/journal.pone.0252431.

Shirvany, Yazdan et al. (2013). “Evaluation of a finite-element reciprocity method for epileptic EEG source localization: Accuracy, computational complexity and noise robustness”. In: Biomedical Engineering Letters 3.1, pp. 8–16. issn: 2093-985X. doi: 10.1007/s13534-013-0083-1.

Tadel, François et al. (2011). “Brainstorm: A User-Friendly Application for MEG/EEG Analysis”. In: Comput. Intell. and Neurosci. 2011.1, p. 879716. doi: 10.1155/2011/879716.

Vallaghé, Sylvain et al. (2008). “The adjoint method for general EEG and MEG sensor-based lead field equations”. In: Phys. Med. Biol. 54.1, p. 135. doi: 10.1088/0031-9155/54/1/009.

Vanrumste, Bart et al. (2001). “The Validation of the Finite Difference Method and Reciprocity for Solving the Inverse Problem in EEG Dipole Source Analysis”. In: Brain Topography 14.2, pp. 83–92. issn: 1573-6792. doi: 10.1023/A:1012909511833.

Wagner, S. et al. (2016). “Using reciprocity for relating the simulation of transcranial current stimulation to the EEG forward problem”. In: NeuroImage 140. Transcranial electric stimulation (tES) and Neuroimaging, pp. 163–173. issn: 1053-8119. doi: 10.1016/j.neuroimage.2016.04.005. URL: https://www.sciencedirect.com/science/article/pii/S1053811916300386.

Wartman, William A et al. (2024). “An adaptive h-refinement method for the boundary element fast multipole method for quasi-static electromagnetic modeling”. In: Phys. Med. Biol. 69.5, p. 055030. doi: 10.1088/1361-6560/ad2638.

Wartman, William A. et al. (2022). “High-Resolution EEG Source Reconstruction with Boundary Element Fast Multipole Method Using Reciprocity Principle and TES Forward Model Matrix”. In: bioRxiv. doi: 10.1101/2022.10.30.514418. URL: https://www.biorxiv.org/content/early/2022/11/01/2022.10.30.514418.

Weise, Konstantin et al. (2022). “The effect of meninges on the electric fields in TES and TMS. Numerical modeling with adaptive mesh refinement”. In: Brain Stimulation: Basic, Translational, and Clinical Research in Neuromodulation 15.3, pp. 654–663. issn: 1935-861X. doi: 10.1016/j.brs.2022.04.009.

Welch, P. (1967). “The use of fast Fourier transform for the estimation of power spectra: A method based on time averaging over short, modified periodograms”. In: IEEE Trans. Audio Electroacoust. 15.2, pp. 70–73. doi: 10.1109/TAU.1967.1161901.

Zarei, Asghar and Babak Mohammadzadeh Asl (2022). “Automatic detection of code-modulated visual evoked potentials using novel covariance estimators and short-time EEG signals”. In: Computers in Biology and Medicine 147, p. 105771. issn: 0010-4825. doi: 10.1016/j.compbiomed.2022.105771. URL: https://www.sciencedirect.com/science/article/pii/S0010482522005431.

Zhang, Zhi (1995). “A fast method to compute surface potentials generated by dipoles within multilayer anisotropic spheres”. In: Phys. Med. Biol. 40.3, p. 335. doi: 10.1088/0031-9155/40/3/001.

Zhu, Yuanlu et al. (2020). “A Hybrid BCI Based on SSVEP and EOG for Robotic Arm Control”. In: Frontiers in Neurorobotics Volume 14 -2020. issn: 1662-5218. doi: 10.3389/fnbot.2020.583641. URL:https://www.frontiersin.org/journals/neurorobotics/articles/10.3389/fnbot.2020.583641.

